# Hypoxia-Sensing CAR T-Cells Provide Safety and Efficacy in Treating Solid Tumors

**DOI:** 10.1101/2020.05.13.091546

**Authors:** Paris Kosti, James W. Opzoomer, Karen I. Larios-Martinez, Rhonda Henley-Smith, Cheryl L. Scudamore, Mary Okesola, Mustafa Y.M. Taher, David M. Davies, Tamara Muliaditan, Daniel Larcombe-Young, Natalie Woodman, Cheryl E. Gillett, Selvam Thavaraj, John Maher, James N. Arnold

## Abstract

There has been significant interest in the prospects of chimeric antigen receptor (CAR) T-cell therapy in the treatment of solid malignancies, and multiple clinical trials are in progress^1^. However, the scope of these trials has been restricted by the lack of availability of tumorspecific targets to direct CAR binding. Tumor specificity is crucial as on-target off-tumor activation of CAR T-cells in healthy tissues can result in potentially lethal toxicities due to uncontrolled cytokine release syndrome^2^. Here we engineer a stringent hypoxia-sensing CAR T-cell system which achieves selective expression of a pan-ErbB-targeted CAR within a solid tumor, a microenvironment characterized by an inadequate oxygen supply. Using murine xenograft models, we demonstrate that despite widespread expression of ErbB receptors in healthy organs, the approach provides anti-tumor efficacy without off-tumor toxicity. This dynamic on/off oxygen-sensing safety switch has the potential to facilitate the unlimited expansion of the CAR T-cell target repertoire for treating solid malignancies.

---

The ErbB family of receptors are an attractive CAR target due to their widespread expression in a range of cancers^3,4^. However, intravenous (i.v.) infusion of anti-ErbB2 CAR T-cells in a patient with metastatic colon cancer resulted in a lethal ‘cytokine storm’ resulting from off-tumor activation of these cells engaging with ErbB2 receptors expressed in the lungs^2^. To investigate this issue, we utilized a 2^nd^ generation pan-anti-ErbB CAR T1 E28z, named T4-CAR, which has specificity towards 8/9 of the possible ErbB receptor homo- and hetero-dimers and binds both human and mouse receptors equivalently^5^. T4-CAR also co-expresses a chimeric cytokine receptor (4αβ) which delivers an intracellular IL-2/IL-15 signal upon binding of IL-4 to the extracellular domain to facilitate *ex vivo* expansion of transduced cells^6–8^ (Fig. 1a). T4-CAR is currently undergoing Phase I evaluation by intra-tumoral (i.t.) delivery in patients with head and neck squamous cell carcinoma (HNSCC; NCT01818323)^9^. Although i.t. delivery of T4-CAR T-cells has proven safe in man, i.v. infusion is desirable as this permits the targeting of both the primary tumor and metastases. I.v. infusion of human T4-CAR T-cells into immunocompromised NSG mice bearing HN3 tumors (Fig. 1b,c and Supplementary Fig. 1a) which express ErbB1-4 (Supplementary Fig. 1b,c) resulted in a lethal toxicity, evident by a rapid loss of weight in these animals (Fig. 1d). As observed clinically^2^, analysis of the blood of these mice revealed evidence of cytokine storm indicated by an increase in pro-inflammatory cytokines (Fig. 1e). In an attempt to resolve the biodistribution of CAR T-cells, we modified the CAR construct to concurrently express a luciferase (Luc) reporter to permit *in vivo* tracking of transduced T-cells (Fig. 1c and Supplementary Fig. 2a). Imaging analysis four days post i.v. infusion of a sub-lethal dose of reporter human CAR T-cells revealed that the majority were accumulating in the lungs and liver, while only a minority were present in the tumor despite the expression of ErbB1-4 (CAR targets) on the tumor cells (Fig. 1f,g and Supplementary Fig. 2b,c). The accumulation in the liver and lungs was not an artefact of the xenograft system as when murine T-cells were transduced to express the human reporter CAR (Extended Data Fig. 1a), these cells also accumulated in these tissues (as well as in the spleen) when infused i.v. into *Rag2^-/-^* mice (Extended Data Fig. 1b,c). Notably, murine T-cell accumulation in the liver, but not the lungs, was CAR-dependent as T-cells expressing the Luc reporter alone were significantly less prevalent at this location (Extended Data Fig. 1c). The CAR-independent T-cell accumulation that was observed in the lungs was likely due to an integrin-dependent interaction as previously described^10^. Profiling of *ErbB1-4* mRNA expression confirmed that all four receptors were expressed in all vital organs, including the lungs and liver (Supplementary Fig. 3a-d). To investigate for direct evidence of T4-CAR T-cell-mediated tissue damage, a sub-lethal dose of human T4-CAR T-cells was infused i.v. into NSG mice and a pathohistological examination using hematoxylin and eosin (H&E) stained tissue sections of the liver and lungs was conducted after 5 days. This analysis revealed the presence of myeloid cell infiltrates, resulting from CAR-mediated inflammation, in the lungs and liver (Fig. 1h,i and Supplementary Fig. 3e). The infiltrate was observed both in a perivascular distribution and scattered throughout the parenchyma, consisting of both neutrophils (polymorphonuclear cells) and large mononuclear cells with abundant cytoplasm, likely to be macrophages (Supplementary Table 1). Hepatocyte necrosis/apoptosis was also seen in some animals (Supplementary Table 1). T4-CAR T-cells accumulated in the kidneys at a lower level (Fig. 1g) with no significant evidence of inflammation in this tissue (Supplementary Table 1). These data indicate that the liver and lungs represent the two key organs for off-tumor CAR T-cell activation.

**Fig. 1.**
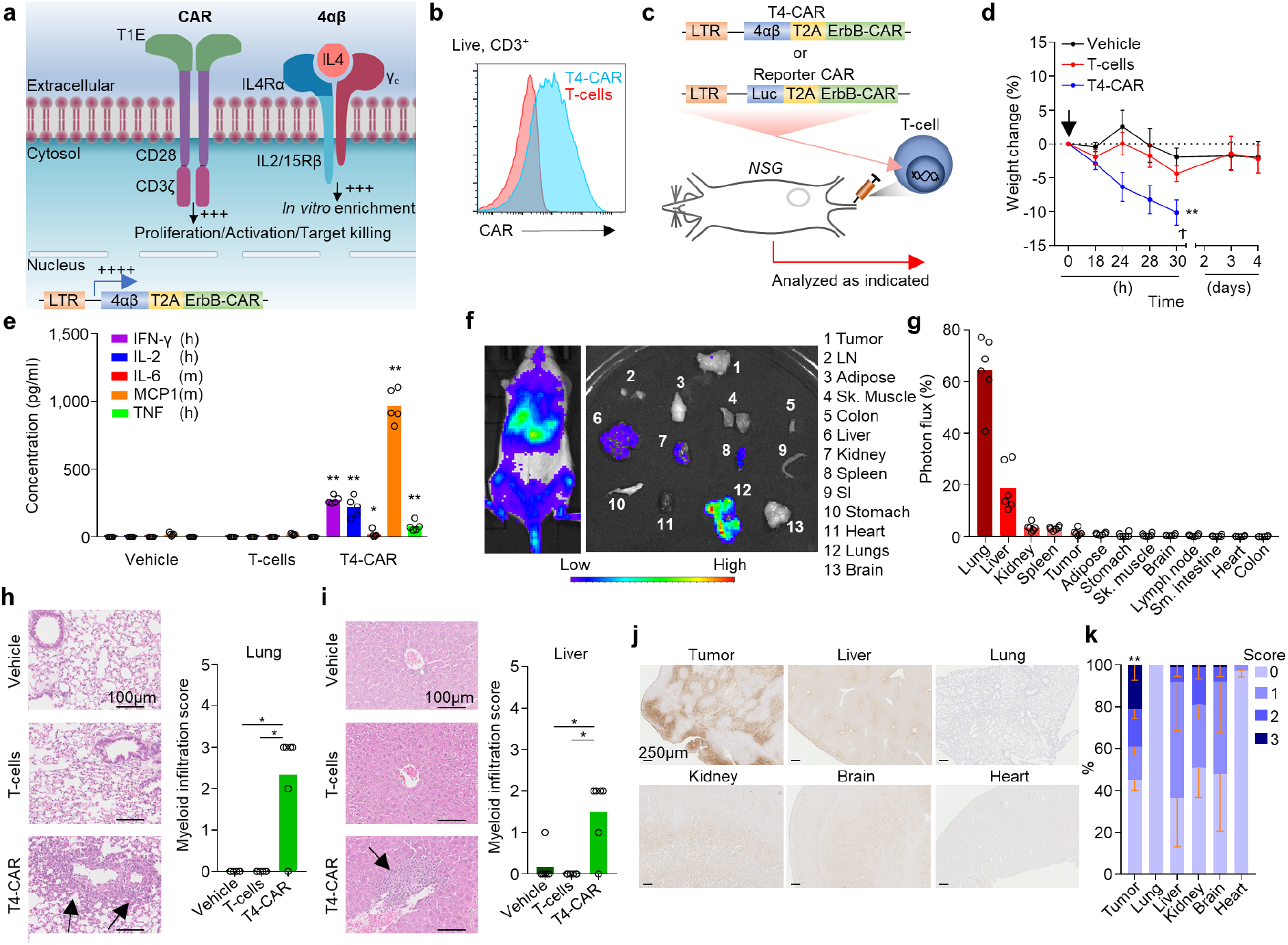
T4-CAR T-cells cause inflammation in healthy organs. (**a**) Diagram depicting T4-CAR. (**b**) Example histograms of surface CAR expression on live (7AAD^-^) CD3^+^ T4-CAR or non-transduced human T-cells assessed using flow cytometry. (**c-e**) Day 13 post subcutaneous HN3 tumor cell inoculation, mice were infused i.v. with vehicle or 10×10^6^ nontransduced or T4-CAR T-cells (n=5). (**c**) Schematic diagram depicting the experiment. (**d**) Weight change of the mice. Arrow denotes T-cell infusion and cross denotes humane endpoint. (**e**) Serum cytokines 24h post-infusion. (**f**) Low-dose human ErbB-CAR/Luc T-cells (4.5×10^6^) were infused i.v. into SKOV3 tumor bearing NSG mice and 4 days later, bioluminescence imaging was performed on the whole body and dissected organs. (**g**) Quantification of the photons/s/unit area as percent of all organs (n=6), LN-inguinal lymph node, SI-small intestine. (**h,i**) H&E stained sections (left) and quantitation of myeloid infiltration (right) in the lung (**h**) and liver (**i**) 5 days post i.v. infusion of low-dose 4.5×10^6^ T4-CAR, non-transduced T-cells or vehicle. Arrows indicate myeloid infiltrates. (**j,k**) IHC staining of tissue sections for reductively-activated pimonidazole in tumor bearing NSG mice (**j**) and quantitation of the staining, scoring between 0-3, from no staining (0) to intense staining (3) as a % area of the tissue (**k**). All experiments are representative of a biological repeat. Line charts, the dots mark mean and error bars s.e.m. Bar charts show mean and dots individual mice. * *P*<0.05, ** *P*<0.01.

Hypoxia is a characteristic of most solid tumors, where proliferative and high metabolic demands of the tumor cells, alongside inefficient tumor vasculature, result in a state of inadequate oxygen supply (<2% O_2_) compared to that of healthy organs/tissues (5-10% O_2_) (Fig. 1j,k and Supplementary Fig. 4)^11^. Clinically, hypoxia has been associated with poor prognosis^12^, and resistance to both chemotherapy^13^ and radiotherapy^14^. Cells have evolved an elegant biological machinery to both detect and rapidly respond to hypoxia through the constitutively expressed transcription factor hypoxia inducible factor alpha (HIF1α)^15^. Under conditions of sufficient O_2_, HIF1α is degraded through hydroxylation of two prolines in an Oxygen-Dependent Degradation Domain (ODD) within its structure^16^. Hydroxylated ODDs are subsequently recognized by von Hippel–Lindau tumor suppressor, which forms part of an E3 ubiquitin ligase complex, that ubiquitinates HIF1α and thereby targets it for proteasomal degradation^17^. Conversely, under limiting O_2_ concentrations HIF1α becomes stabilized and translocates to the nucleus where it binds to HIF1β and p300/CBP. This complex can then associate with Hypoxia Responsive Elements (HREs) in the promoter region of hypoxia-responsive genes initiating transcription^18^. As hypoxia differentiates the tumor microenvironment from that of healthy, normoxic tissue, it represents a desirable marker for the induction of CAR T-cell expression (Fig. 1j,k and Supplementary Fig. 4). A previous study has investigated CARs fused with an ODD^19^. However, although this approach endowed CAR T-cells with an improved ability to kill tumor cells under hypoxic conditions *in vitro,* the authors observed residual tumor cell killing under normoxic conditions. In an attempt to create a stringent hypoxia-regulated CAR expression system, we developed a dual-oxygen sensing approach for the T4-CAR (Fig. 2a). This was achieved by appending a C-terminal 203 amino acid ODD^20^ onto the CAR while concurrently modifying the CAR’s promoter in the long terminal repeat (LTR) enhancer region of the retroviral vector to contain a series of 9 consecutive HREs. This CAR, named ‘HypoxiCAR’, *in vitro* demonstrated stringent hypoxia-specific presentation of the CAR molecules on the cell surface of human T-cells (Fig. 2b). The dual-hypoxia sensing system incorporated into HypoxiCAR proved superior to either single-hypoxia sensing modules of the 9xHRE cassette or ODD, which displayed leakiness in CAR expression (Extended Data Fig. 2a,b) and tumor cell killing (Extended Data Fig. 2c) under conditions of normoxia. HypoxiCAR’s expression of the CAR, utilizing the dual-hypoxia sensing system, was both stringently regulated to hypoxic environments and was also highly dynamic, representing a switch that could be turned both ‘on’ and ‘off’ in an O_2_-dependent manner (Fig. 2c and Extended Data Fig. 2b). As the 4αβ receptor of T4-CAR was not appended to an ODD, the leaky expression of the 9xHRE promoter under conditions of normoxia (Extended Data Fig. 2b) was sufficient to allow IL-4 mediated *in vitro* expansion of HypoxiCAR T-cells under culture conditions of normoxia (Fig. 2a). In further *in vitro* characterization, the exquisite O_2_ sensitivity of HypoxiCAR was confirmed where CAR expression was absent under O_2_ concentrations consistent with healthy organs (≥5%) but became detectable on the cell surface at O_2_ concentrations equivalent to those found in the tumor microenvironment (≤1%) (Fig. 2d).

**Fig. 2.**
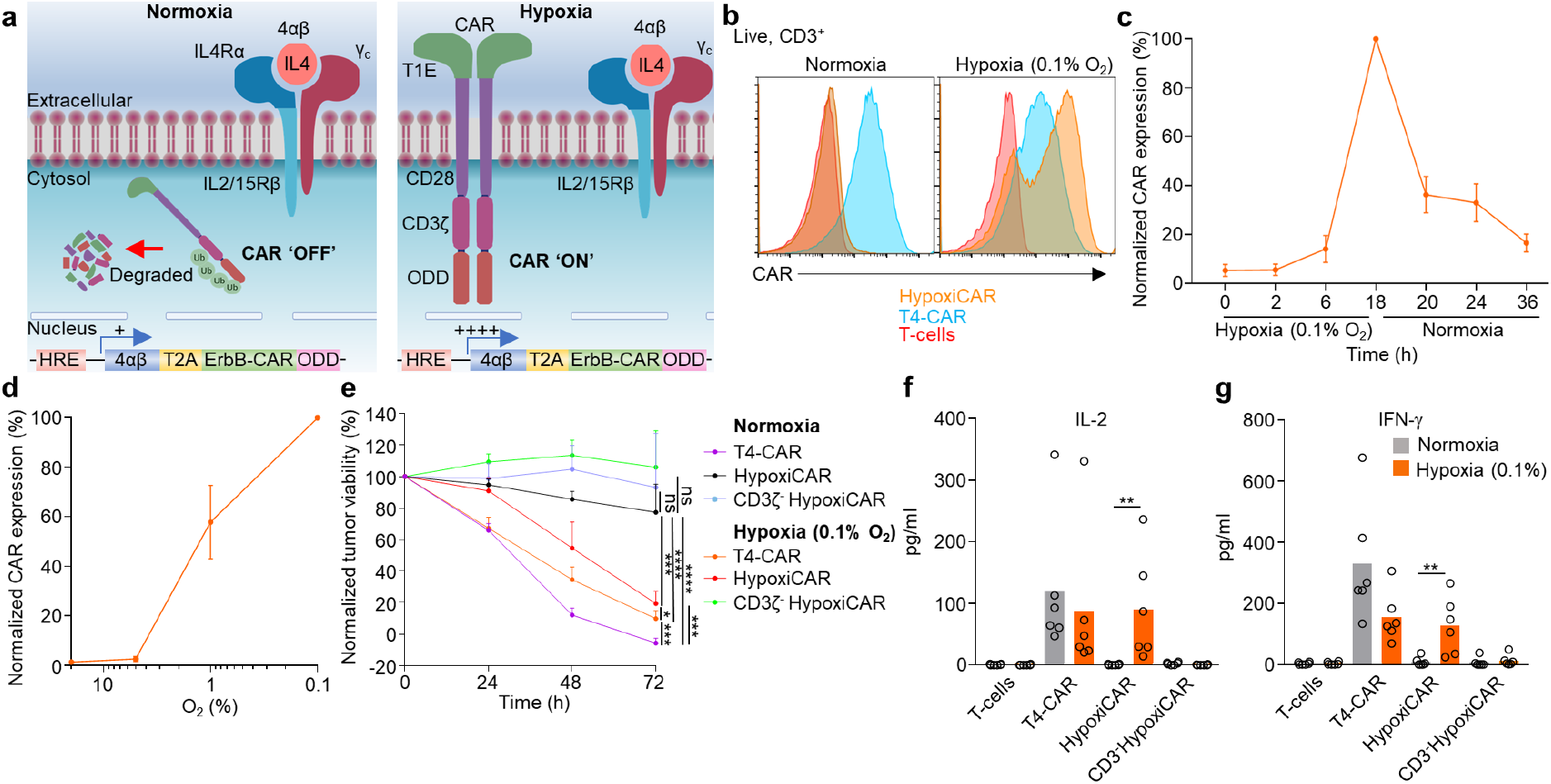
HypoxiCAR T-cell effector function is stringently restricted to hypoxia. (**a**) Diagram depicting HypoxiCAR in normoxia and hypoxia. (**b**) Example histograms of surface CAR expression on live (7AAD^-^) CD3^+^ T4-CAR, HypoxiCAR and non-transduced human T-cells in normoxic or 18h hypoxic (0.1% O_2_) conditions assessed using flow cytometry. (**c**) Surface CAR expression on HypoxiCAR at the indicated times under hypoxia (0.1% O_2_) or normoxia assessed using flow cytometry analysis, values normalized to 18h hypoxia (n=6). (**d**) Surface CAR expression on HypoxiCAR T-cells after 18h exposure to 0.1, 1, 5%, 20% O_2_ (n=6), values normalized to 0.1% O_2_. (**e-g**) *In vitro* SKOV3 tumor cell killing by T4-CAR, HypoxiCAR, CD3ζ-truncated HypoxiCAR (CD3ζ-; to prevent intracellular signaling) and nontransduced T-cells (CAR^+^ T-cell effector to target tumor cell ratio 1:1) in normoxic and 0.1% O_2_ hypoxic conditions (**e**). Quantification of IL-2 (**f**) and IFN-γ (**g**) released into the media from the respective T-cells after 24h and 48h exposure to SKOV3 cells respectively, under normoxic and 0.1% O_2_ hypoxic conditions. Bar on charts shows mean and dots an individual healthy donor, datapoints collected in parallel and representative of a biological repeat. In line charts, the dots mark mean and error bars s.e.m. * *P*<0.05, ** *P*<0.01, *** *P*<0.001, **** *P*<0.0001.

Having validated HypoxiCAR’s ability to sense hypoxia, we sought to investigate its ability to elicit hypoxia-dependent killing of tumor target cells. SKOV3 ovarian cancer cells were seeded onto culture plates and co-incubated with T4-CAR or HypoxiCAR under normoxic and hypoxic (0.1% O_2_) conditions. Despite equivalent transduction efficiencies and CD4^+^:CD8^+^ T-cells ratios (Supplementary Fig. 5), HypoxiCAR T-cells displayed efficient hypoxia-dependent killing of the SKOV3 cells, almost equivalent to T4-CAR T-cells, with no significant killing observed under normoxic conditions (Fig. 2e). Target-cell destruction was strictly CAR-dependent as when the intracellular tail of HypoxiCAR was truncated, to prevent CD3ζ signaling, killing was abrogated (Fig. 2e). In addition, HypoxiCAR-provided stringent hypoxia-restricted T-cell secretion of both IL-2 (Fig. 2f) and IFN-γ (Fig. 2g), two cytokines which play an important role in the T-cell response^21,22^.

To evaluate whether HypoxiCAR could provide tumor-restricted CAR expression *in vivo,* human HypoxiCAR T-cells were injected concurrently i.v. and i.t. in NSG mice bearing HN3 tumors with an approximate volume of 500mm^3^ (Fig. 3a and Supplementary Fig. 6a). The presence of hypoxia in tumors of this volume was confirmed (Fig. 1j,k and Supplementary Fig. 4). Four days after HypoxiCAR T-cell infusion, tissues were harvested, enzyme-digested and T-cells were assessed for CAR expression using flow cytometry (Supplementary Fig. 6b). As predicted by the *in vitro* analyses (Fig. 2), HypoxiCAR T-cells had no detectable surface CAR molecules when recovered from the blood, lungs, or liver of the mice, but they did express surface CAR molecules within the hypoxic tumor microenvironment (Fig. 3b,c). This finding was not model specific as similar observations were made in both NSG mice bearing SKOV3 tumors (Extended Data Fig. 3a,b) and in *Rag2^-/-^* mice bearing murine Lewis lung carcinoma (LL2) tumors (Extended Data Fig. 3c).

**Fig. 3.**
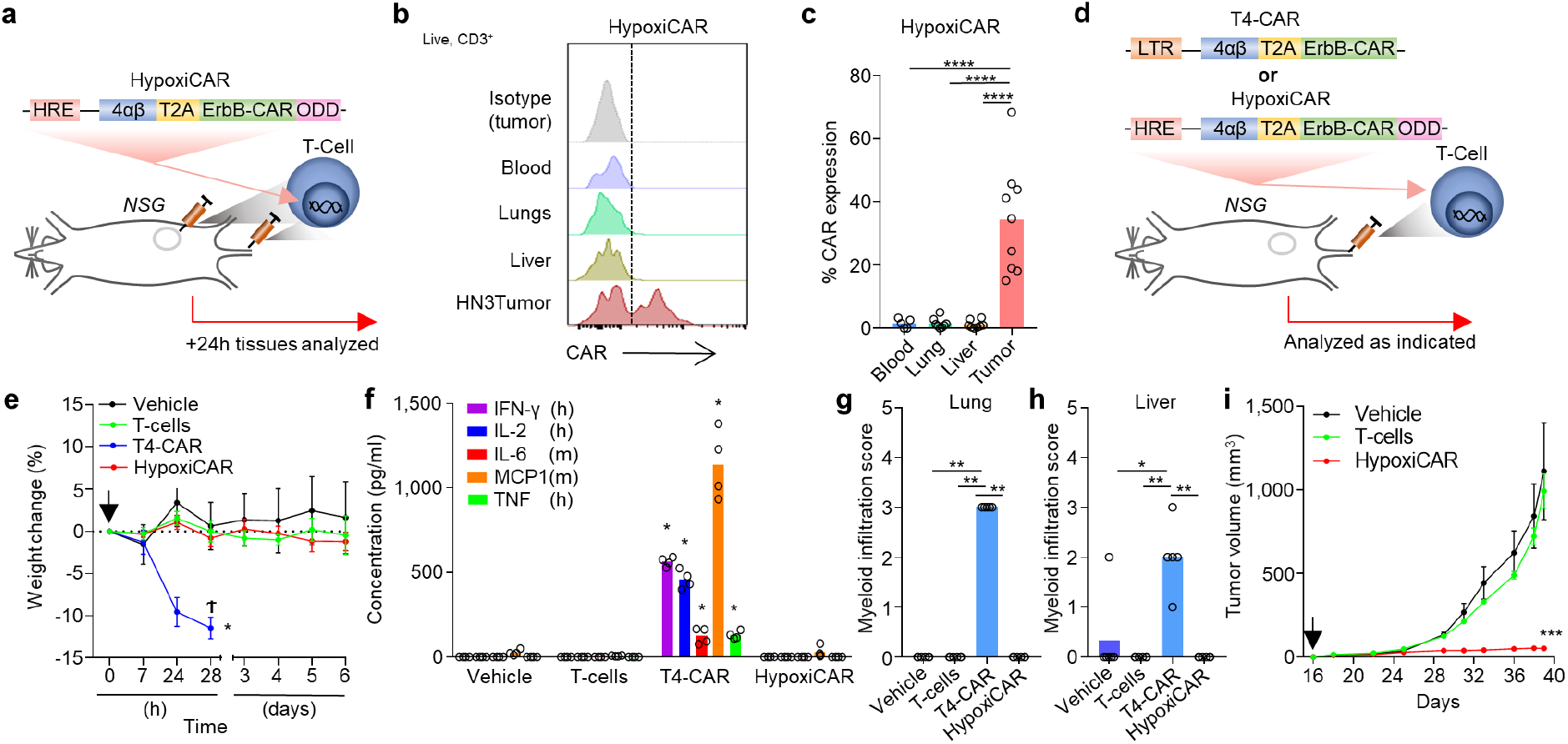
HypoxiCAR T-cells provide anti-tumor efficacy without systemic toxicity. (**a-c**) Subcutaneous HN3 tumor-bearing NSG mice were injected both i.v. and i.t. with human HypoxiCAR T-cells (2.5×10^5^ i.t. and 7.5×10^5^ i.v.) 72h prior to sacrifice. (**a**) Schematic diagram depicting the experiment. (**b**) Representative histograms showing surface CAR expression on live nucleated cells (7AAD^-^, Ter119^-^), CD45^+^ CD3^+^ HypoxiCAR T-cells in the indicated enzyme-dispersed tissues and blood and (**c**) quantification in the respective tissues across n=9 individual mice. (**d-f**) Sixteen days post subcutaneous HN3 tumor cell inoculation, mice were infused i.v. with either vehicle or 10×10^6^ T4-CAR, HypoxiCAR or nontransduced human T-cells (control) (n=4 mice). (**d**) Schematic diagram depicting the experiment. (**e**) Weight change of the mice. (**f**) Serum cytokines 24h post-infusion. (**g,h**) Low dose (4.5×10^6^) T4-CAR or HypoxiCAR T-cells were infused i.v. into NSG mice (n=5-6). Five days later the indicated tissues were excised, and myeloid infiltration was scored in the lung (**g**) and liver (**h**). (**i**) HN3 tumor growth curves from (**d-f**), arrow marking the point of CAR T-cell infusion. All experiments are representative of a biological repeat. Bar charts shows the mean and each dot an individual mouse. In line charts, the dots mark the mean and error bars s.e.m. * *P*<0.05, ** *P*<0.01, *** *P*<0.001, **** *P*<0.0001.

To test the anti-tumor efficacy of HypoxiCAR, high-dose T4-CAR, HypoxiCAR or nontransduced human T-cells were infused into mice at day 16 post injection of HN3 tumor cells. In keeping with the absence of CAR expression on the T-cells in normoxic tissues (Fig. 3b), HypoxiCAR circumvented the treatment-limiting toxicity seen following i.v. infusion of high-dose T4-CAR T-cells. Mice infused with HypoxiCAR T-cells displayed no acute drop in weight post-infusion (Fig. 3d,e), no evidence of pro-inflammatory cytokines in the systemic circulation (Fig. 3f), nor signs of tissue damage in the lungs, liver or kidneys (Fig. 3g,h, Supplementary Fig. 7 and Supplementary Table 2). Importantly, while mice infused i.v. with human T4-CAR T-cells all reached their humane endpoints at 28 h (Fig. 3e), those infused with HypoxiCAR T-cells displayed no signs of toxicity while tumor growth was effectively prevented (Fig. 3i). This observation was also not model specific, as HypoxiCAR T-cells also circumvented any toxicity-related weight loss (Extended Data Fig. 4a,b) and suppressed tumor growth in mice bearing established SKOV3 tumors (Extended Data Fig. 4c). To directly confirm that HypoxiCAR T-cells accumulated at the site of disease when tumor control was observed, HypoxiCAR T-cells were co-transduced to express a constitutively expressed *Renilla* (r)Luc. Tracking the biodistribution of HypoxiCAR T-cells *in vivo* confirmed their infiltration into the tumor microenvironment and persistence in these animals for at least 26 days post infusion (Extended Data Fig. 4d-f). As such, HypoxiCAR overcomes a major hurdle that currently precludes the systemic administration of CAR T-cells targeting antigens that are expressed in normal tissues throughout the body^2^.

Hypoxia has been extensively studied in HNSCC^12,23,24^. To assess which patients might be most appropriate for HypoxiCAR T-cell immunotherapy, we firstly generated an HRE-regulated gene signature using patient tumor sample transcriptomic data. Known HRE-regulated genes were analyzed for co-expression, and a refined signature utilizing the genes *PGK1, SLC2A1, CA9, ALDOA* and *VEGFA* was chosen as we observed a significant positive correlation between these genes (Fig. 4a and Supplementary Fig. 8a). There was no difference in expression of this signature across the different HNSCC subtypes (hypopharynx, larynx, oral cavity, and oropharynx; Supplementary Fig. 8b). However, expression of this 5-gene signature, significantly increased with tumor size (T-stage; Fig. 4b) and was also prognostic of poorer survival in stage 3 and 4 HNSCC patients (Fig. 4c). All five HRE-regulated genes within the signature were also upregulated in subcutaneous human tumor cell line xenografts grown in NSG mice, such as were utilized in this study (Supplementary Fig. 8c). Other groups have also demonstrated hypoxia-gene signatures to be predictive of adverse prognosis in HNSCC^25–27^. Such signatures could be utilized to predict hypoxia from tumor biopsy samples as a means to select those patients who might respond best to HypoxiCAR therapy.

**Fig. 4.**
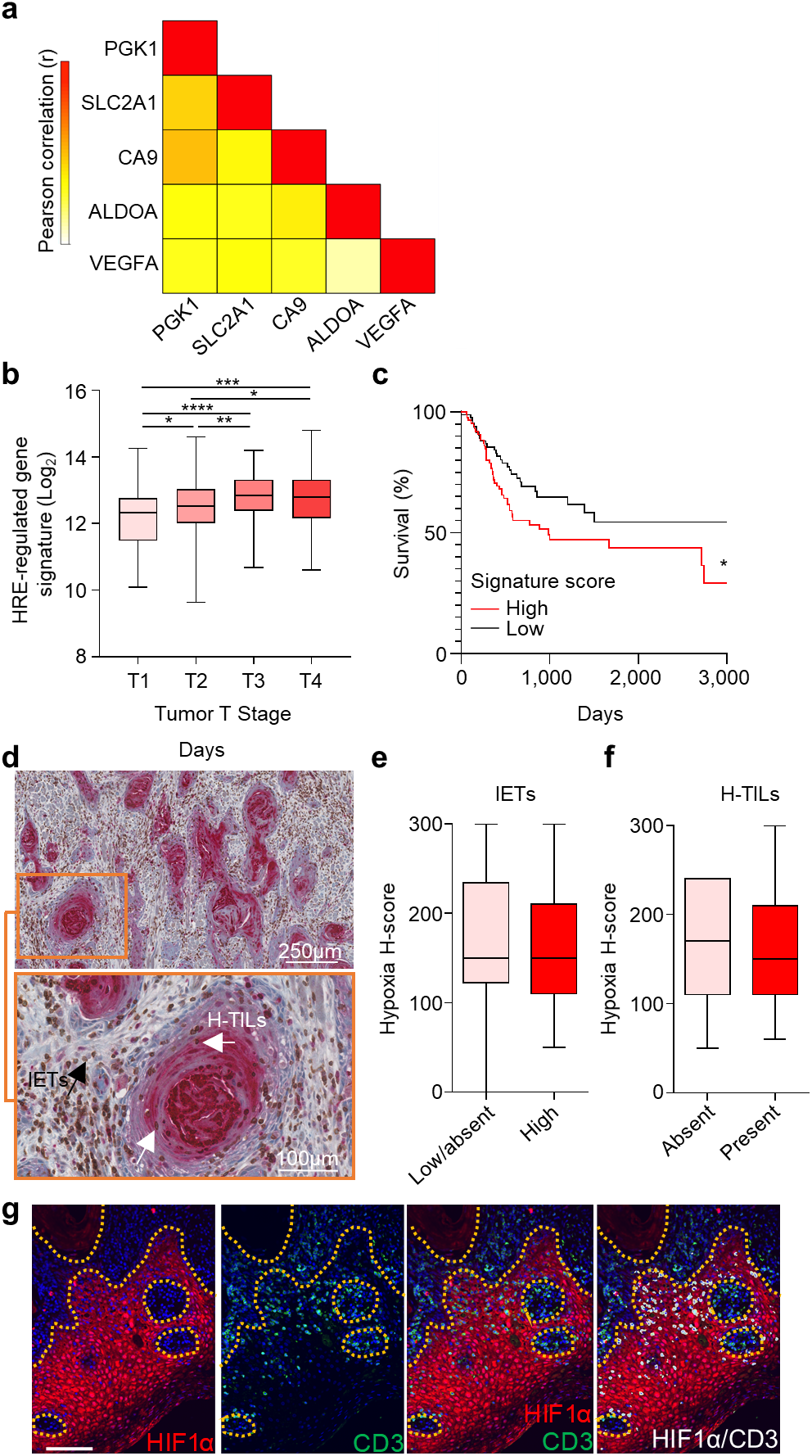
T-cells are not excluded from HIF1α stabilized regions of hypoxic HNSCCs. (**a c**) An HRE-regulated gene signature was constructed from known HRE-regulated genes in the HNSCC TCGA dataset (n=528). (**a**) Heatmap displaying the Pearson correlation coefficient for the individual genes in the TCGA HNSCC dataset. (**b**) Signature expression based on T stage (T1 n=48, T2 n=136, T3 n=99, T4 n=174). (**c**) Survival curve for patients with Stage 3 and 4 HNSCC for high and low expression of the HRE-regulated gene signature (n=87 respectively). (**d**) Example IHC stained HNSCC section for HIF1α (red) and CD3 (brown) (n = 60). (**e-f**) The abundance of inter-epithelial T-cells (IETs), example marked by black arrow (D), low/absent n=40 and high n=55 assessed against the HIF1α stabilization score of the tumor (**e**) and, for the tumors where IETs were high, TILs directly infiltrating HIF-1α stabilized regions of the tumor (H-TILs, examples marked by white arrows (**d**)) as absent (n=6 of 55 tumors) or present (n=46 of 55 tumors) (**f**). (**g**) Confocal images from an oral tongue carcinoma stained with DAPI (nuclei; blue) and antibodies against CD3 (green) and HIF1α (red); white denotes CD3 and HIF1α co-localization. Box plots show median and upper/lower quartiles, whiskers show highest and lowest value. * *P*<0.05, ** *P*<0.01, *** *P*<0.001, **** *P*<0.0001.

Immunohistochemistry staining of HNSCC tumor sections for stabilized HIF1α, the master transcription factor for HypoxiCAR’s CAR expression, revealed large regions of the tumors where HIF1α had become stabilized (Fig. 4d). Although several factors can stabilize HIF1α, hypoxia represents the most probable explanation for this observation^28^. Heterogeneity in both HIF1α stabilization and intra-tumoral T-cell infiltration was seen between patients. Encouragingly however, those tumors with the highest prevalence and/or intensity of HIF1α stabilization did not exclude T-cells from entering the inter-epithelial space (Fig. 4e), nor from entering HIF1α stabilized regions of the tumor (Fig. 4f). Using immunofluorescence, we also demonstrated that CD3^+^ T-cells infiltrating HIF1α stabilized tumor regions also stabilized HIF1α themselves, suggesting that in these environments HypoxiCAR T-cells would become activated (Fig. 4g and Supplementary Fig. 8d). These observations suggest that HypoxiCAR could find clinical application in hypoxic tumor types such as HNSCC, where gene expression (Fig. 4a-c and Supplementary Fig. 8a,b), staining of biopsy samples for HIF1α/CD3 (Fig. 4d-g and Supplementary Fig. 8d) and imaging techniques such as PET/CT using a hypoxia-radiotracer such as ^64^Cu-ATSM^24^ might provide biomarkers to confirm the presence of a hypoxic tumor microenvironment and guide patient selection^29^.

Approaches to improve tumor-specificity of CAR T-cells have been developed, such as T-cell receptor-mimetic CARs with specificity for HLA-presented antigens^30^, combined targeting of tumor antigens^31–33^, or tuning of CAR affinity to preferentially target high density antigens^34^. This study demonstrates an alternative approach to achieve cancer-selective immunotherapy, exploiting one of the most innate characteristics of the tumor microenvironment. The dual ‘hypoxia-sensing’ system described here achieves compelling anti-tumor efficacy while abrogating off-tumor toxicity of a CAR that recognizes multiple targets in normal tissues. ‘HypoxiCAR’ using a pan-ErbB-targeted CAR represents the first incarnation of this approach and provides a stringent and broadly applicable strategy to overcome the paucity of safe targets available for the treatment of solid malignancies.

## Online Methods

### Mice

NSG (NOD-*scid* IL2Rgamma^null^) mice were purchased from Charles River. Balb/c *Rag2^-/-^* mice were a gift from Professor Adrian Hayday (KCL). Male mice were used for studies involving HN3 and female mice were used for studies involving SKOV3 and LL2 studies. All mice used for ectopic tumor studies were 6-8 weeks old and approximately 22 g in weight.

### Cell lines

SKOV3 human ovarian adenocarcinoma cells^35^ were originally purchased from ATCC and were re-authenticated for this study by ATCC. HN3 human head and neck adenocarcinoma cells^5,7^ were acquired from Ludwig Institute for Cancer Research, London and grown in D10 medium, Dulbecco’s modified Eagle’s medium (DMEM; Gibco) supplemented with 10% Fetal calf serum (FCS; Thermo Fisher Scientific) and GlutaMAX (Thermo Fisher Scientific). Murine Lewis Lung carcinoma (LL2)^36^ cells were purchased from ATCC and were cultured in RPMI 1640 (Gibco) supplemented with 10% FCS (Thermo Fisher Scientific). Cell lines were confirmed to be free of mycoplasma for this study using the MycoAlert^®^ Mycoplasma Detection Kit (Lonza).

### CAR/Reporter construct cloning

Human T1 E28z CAR (pan-ErbB-targeting CAR) containing SFG retroviral vector^5,7,37^ was modified to generate the constructs utilized in this study. The full-length ODD cDNA encoding for amino acids 401-603 from human HIF1α was synthesized as a gBlock^®^ (Integrated DNA Technologies) and was appended onto the C-terminus of CD3ζ within the T1E28z CAR through overlap PCR using Platinum *Pfx* DNA polymerase (Thermo Fisher Scientific) according to the manufacturer’s instructions. The forward primer 5’-GCCTACCAAGAACAACTGGAC-3’ and reverse primer 5’-TCCAGCGGCTGGGGCGCGAGGGGGCAGGGCC-3’ were used to amplify 4αβ-T2A T1E28z CAR while introducing ODD-compatible sticky ends. The ODD was amplified using forward primer 5’-GGCCCTGCCCCCTCGCGCCCCAGCCGCTGGA-3’ and reverse primer 5’-GACTAATCCGGATCCTCGAGTGGCTGTTACTGGAATACTGTAACTGTGCTTTGAGG-3’ which also introduced a CD3ζ-compatible sticky end. The PCR fragments were then fused using the forward primer 5’-GCCTACCAAGAACAACTGGAC-3’ and 5’-GACTAATCCGGATCCTCGAGTGGCTGTTACTGGAATACTGTAACTGTGCTTTGAGG-3’. PCR products were run on 1.2% Agarose (Sigma-Aldrich) gels and product size was estimated against a 1kb Plus DNA ladder (Thermo Fisher Scientific). Fragments of the expected size were excised and purified using the QIAquick^®^ Gel Extraction kit. The 4αβ T2A T1 E28z CAR-ODD sequence which contained flanking *AgeI* and *XhoI* cleavage sites was cloned into the SFG vector to replace the wild type T1 E28z CAR with the T1 E28z ODD CAR. AgeI and XhoI restriction endonucleases (New England Biolabs) were used to cleave *AgeI* and *XhoI* restriction enzyme cleavage sites in the SFG plasmid to remove the existing 4αβ T2A CAR from the vector backbone. AgeI and XhoI restriction endonucleases were also used to cleave the *AgeI* and *XhoI* restriction enzyme sites which were built to flank the 4αβ T2A T1E28z CAR-ODD cDNA. Vector and constructs that had been restriction endonuclease digested were purified using QIAquick PCR purification kit (Qiagen) and ligated using T4 ligase (Thermo Fisher Scientific). The final ODD modified 4αβ T2A T1 E28z CAR-ODD construct was 3212bp in length (including the 609bp ODD) from the start to the stop codon. Plasmids were transformed into One Shot Stbl3™ chemically competent *E. coli* (Thermo Fisher Scientific). Transformed *E. coli* were selected using ampicillin (Santa Cruz Biotechnology) containing Luria Bertani (LB) Agar (Sigma-Aldrich) plates. Transformed colonies were then grown up in LB broth (Sigma-Aldrich) with 100 μg/ml ampicillin and purified using either Qiagen Plasmid Midi or Maxi kits. Final constructs were sequence verified (Source BioScience). The constitutively expressed reporter construct has previously been described^38^ and contained a Click Beetle Luciferase (Luc) and eGFP separated by a viral P2A sequence^39^. The reporter construct was PCR amplified using Platinum *Pfx* DNA polymerase (Thermo Fisher Scientific) according to the manufacturer’s protocol with the forward primer 5’-CCATGGTGAAGCGTGAGAAAAATG-3’ and the reverse primer 5’-CTCGAGTTACTTGTACAGCTCGTCCATGC-3’. The amplified product was digested with NcoI and XhoI restriction endonuclease (New England Biolabs) and cloned into the SFG vector using the *Ncol* and *Xhol* restriction sites and T4 DNA ligase (Thermo Fisher Scientific). The HRE modification was targeted in the 3’ LTR of the SFG retroviral vector, as the 3’ LTR region is copied to the 5’ LTR upon integration^40^. DNA containing 9 tandem 5’-GGCCCTACGTGCTGTCTCACACAGCCTGTCTGAC-3’ HRE motifs (total length 306bp) derived from the human *EPO* gene, contained both HIF-binding and ancillary sites, was synthesized as a gBlock^®^ (Integrated DNA Technologies) and sub-cloned into the 3’ LTR of the SFG vector, to replace an equivalent sized fragment within the enhancer region, between the *NheI* and *XbaI* restriction endonuclease sites upstream of the native murine leukemia virus promoter. As *NheI* and *XbaI* were not unique restriction sites in the vector, to achieve specific modification of the 3’ LTR, we generated using overlapping fusion PCR a larger fragment flanked *XhoI/EcoRI* unique restriction endonuclease sites. This fragment was identical to the vector DNA *XhoI/EcoRI* region except that the *NheI/XbaI* region was replaced by the 9 HRE gBlock^®^. The T1E28z CAR CD3^-^ truncated control construct was synthesized as gBlock^®^ (Integrated DNA Technologies) with flanking *SbfI* and *XhoI* restriction sites and sub-cloned into the HRE-modified SFG vector using SbfI and XhoI restriction endonucleases (New England Biolabs). To generate the bicistronic Luciferase-T2A-CAR construct for *in vivo* tracking CAR T-cells, a gBlock^®^ (Integrated DNA Technologies), which was designed to include Luciferase-T2A-T1E peptide binder flanked with *AgeI* and *NotI* restriction sites, was inserted into the T1E28z CAR construct.

### T-cell isolation

For isolating murine T-cells; spleens were excised from WT C57Bl/6 mice and placed in complete media which contained RPMI 1640, 10% FCS, 20 μM 2-mercaptoethanol, 1X penicillin/streptomycin (Sigma-Aldrich). Spleens were crushed then washed through a 70 μm pore strainer and RPMI 1640. Liberated splenocytes were centrifuged at 500 *g* for 5 mins and the cell pellet was re-suspended in 1 ml of red blood cell lysis buffer (Roche) for 2 mins at RT. Cells were then centrifuged again at 500 *g* for 5 mins and the pellet resuspended in RPMI 1640 and then cells were numerated using Trypan blue (Sigma-Aldrich) exclusion on a hemocytometer. CD3^+^ T-cells were purified using the murine Pan T-cell Isolation Kit II (Miltenyi Biotec) and isolated using a MidiMacs separator and LS columns (Miltenyi Biotec) according to the manufacturer’s protocol. T-cells were resuspended in complete media supplemented with 2 ng/ml recombinant murine IL-2 (Bio-Techne) and plated at a density of 0.1×10^6^ cells/well in 200 μl onto a high binding 96-well plate (Sigma-Aldrich) that had been pre-coated overnight with a mix of anti-mouse CD3ε (145-2C11,5 μg/ml) and anti-mouse CD28 (37.51,3 μg/ml) antibodies in sterile PBS (100 μl/well) at 4°C.

For isolating human T-cells; blood was obtained from healthy volunteers under approval of the Guy’s and St Thomas’ Research Ethics Committee (REC reference 09/H0804/92). Blood was collected into Falcon tubes containing anti-coagulant (10% Citrate), mixed at 1:1 with RPMI 1640 and layered over Ficoll-Paque Plus (GE Healthcare). Samples were centrifuged at 750 *g* for 30 mins at 20°C (acceleration and brake set to 0) to separate the peripheral blood mononuclear (PBMC) cell fraction. The interface between the plasma and the Ficoll layer, which contained the PBMCs, was harvested using a sterile Pasteur pipette and washed in RPMI 1640. T-cells were purified from the PBMC fraction using human Pan T-cell isolation kit (Miltenyi Biotec) and isolated using a MidiMACs™ separator and LS columns (Miltenyi Biotec) according to the manufacturer’s protocol. Purified human T-cells were activated using CD3/CD28 Human T-Activator Dynabeads (Gibco) at a 1:1 cell to bead ratio and seeded in tissue culture plates at 3×10^6^/ml in RPMI 1640 supplemented with 5% human serum (Sigma-Aldrich) and 1X penicillin/streptomycin. The following day, 100 IU/ml recombinant human IL-2 (PROLEUKIN) was added to the cultures.

### Retroviral transduction

To produce retrovirus with tropism for human cells, RD114 pseudotyped retroviral particles were generated by triple transfection, using Peq-Pam plasmid (Moloney GagPol; a gift from Dr Martin Pule, UCL), RDF plasmid (RD114 envelope; a gift from Prof. Mary Collins, UCL) and the SFG plasmid of interest, using FuGENE HD transfection reagent (Promega), of HEK 293T cells as previously described^41^. For the *in vivo* experiments evaluating the therapeutic efficacy of CAR T-cells in which the SKOV3 tumor cells expressed Click Beetle luciferase, T-cells were tracked by co-transduction with a construct containing *Renilla* luciferase. Cotransduction was conducted using 1:1 ratios of virus containing a red-shifted *Renilla* luciferase 8.6-535 variant (rluc; Genscript)^42^, which contained the *Renilla* luciferase separated from eGFP by a furin-T2A sequence (rluc/eGFP)^43^, alongside the respective retroviral particles containing the indicated CAR constructs. To produce retrovirus with murine cell tropism, the Phoenix-ECO retrovirus producer cell line was transfected using FuGENE HD (Promega) with the relevant plasmid. Supernatant containing viral particles were harvested and incubated with the cells of interest for at least 48 h to allow their transduction. T-cells were transduced in non-tissue culture treated plates that were precoated with 4 μg/cm^2^ RetroNectin (Takara Bio) overnight at 4°C. Prior to the retroviral transduction of human T-cells, CD3/CD28 Human T-Activator Dynabeads (Gibco) were removed and fresh IL-2 was added as stated in the T-cell isolation section. In the case of T-cell transduction with the bicistronic 4αβ-T2A-CAR construct, following T-cell transduction, human IL-4 (Peprotech) at 30 ng/ml final concentration was added to the culture to selectively enrich the transduced T-cell population.

### Quantitative PCR

Genomic DNA was extracted from cells using a DNeasy Blood & Tissue Kit (Qiagen) according to manufacturer’s protocol and quantitative PCR was performed using KiCqStart SYBR Green qPCR ReadyMix with ROX (Sigma-Aldrich) according to the manufacturer’sprotocol using custom designed primers to generate amplicons from *Tbp, Luc* or *T2A* sequences in the genome. The primers used were: murine *Tbp* 5’-TGTCTGTCGCAGTAAGAATGGA-3’ and 5’-AAAATCCCAGACACGGTGGG-3’, human *TBP* 5’-TTTGGTGTTTGCTTCAGTCAG-3’ and 5’-ATACCTAGAAAACAGGAGTTGCTCA-3’, *Luc* 5’-ATTTGACTGCCGGCGAAATG-3’ and 5’-AAGATTCATCGCCGACCACAT-3’, *T2A* 5’-CGGAGAAAGCGCAGC-3’ and 5’-GGGTCCGGGGTTCTCTT-3’. Amplification of the genes of interest was detected on an ABI 7900HT Fast Real Time PCR instrument (Thermo Fisher Scientific).

### Quantitative reverse transcriptase PCR

mRNA was extracted using TRIzol (Thermo Fisher Scientific) method and quantitative reverse transcription (qRT) PCR was performed as previously described^44^ using the EXPRESS one-step Superscript RT PCR kit and the following primers/probes purchased from Thermo Fisher Scientific: *ALDOA* Hs00605108_g1, *CA9* Hs00154208_m1, *Erbb1* Mm01187858_m1, *Erbb2* Mm00658541_m1, *Erbb3* Mm01159999_m1, *Erbb4* Mm01256793_m1, *PGK1* Hs99999906_m1, *SLC2A1* Hs00892681_m1, *TBP* Hs00427620_m1, *Tbp* Mm01277045_m1 and *VEGFA* Hs00900055_m1. Expression of all genes is represented relative to the house-keeping gene Tata-binding protein for both human and murine experiments. Assays were performed using an ABI 7900HT Fast Real Time PCR instrument (Thermo Fisher Scientific).

### *In vitro* studies

*In vitro* hypoxia was achieved using a hypoxia incubator chamber (Stemcell Technologies) purged at 25L/min for 4 mins with gas containing either; 0.1, 1 or 5% O_2_, 5% CO_2_ and nitrogen as a balance (BOC), after which the chamber was sealed. This process was repeated again after 1 h. In *in vitro* cytotoxicity assays 1×10^4^ Luc/eGFP-expressing SKOV3 cells were seeded in 96-well tissue culture plates and transduced or non-transduced T-cells were added in the well at the indicated effector to target ratios. Co-cultures were incubated for 24, 48 and 72 h time points in normoxia or experimental hypoxia as indicated and target cell viability was determined by luciferase quantification following the addition of 1 μl of 15mg/ml XenoLight D-luciferin (PerkinElmer) in PBS per 100μl of media. Luminescence was quantified using a FLUOstar Omega plate reader (BMG Labtech). At the 24 and 48 h coculture time points a sample of media was taken from the co-culture and subsequently used for IL-2 and IFN-γ quantification, respectively. IL-2 was quantified using Human IL-2 ELISA Ready-SET-Go! Kit, 2nd Generation (eBioscience) as per manufacturer’s protocol. IFN-γ was quantified using Human IFN-γ DuoSet ELISA kit (Bio-Techne) as per manufacturer’s protocol. In both ELISAs cytokine concentration was determined by absorbance measurements at 450 nm on a Fusion alpha-FP spectrophotometer (PerkinElmer).

### *In vivo* studies

Tumor cell lines (2.5×10^5^ cells in PBS) were inoculated by subcutaneous (s.c.) injection into female (for SKOV3 and LL2) and male (for HN3) mice that were six to eight weeks of age. Once tumors were palpable, digital caliper measurements of the long (L) and short (S) dimensions of the tumor were performed every 2 or 3 days. Tumor volume was established using the following equation: Volume= (S^2^xL)/2. The indicated doses of CAR T-cells were injected in 200μl PBS through the tail vein using a 26 G needle. Where i.t. injection was used, cells were injected directly into the tumor in 50μl PBS. Blood samples were taken from mice in EDTA-coated Microvette™ tubes (Sarstedt) and plasma was extracted by centrifugation of these samples at 2,000 *g* for 5 mins. Cytokine concentrations were measured externally by Abcam using the FirePlex™ Human Th1/Th2/Th17 and FirePlex™ Mouse Inflammation Immunoassay panels. After mice had been humanely sacrificed at the end of a study period, where tissues were excised they were immersed in excess formalin solution (10%) neutral buffered (Sigma-Aldrich), paraffin embedded, sectioned and stained with H&E using standard protocols. Blinded analysis of histopathology was performed by a FRCPath-qualified specialist in veterinary pathology. Scores were assigned for evidence of inflammation using a non-linear semi-quantitative grading system from 0 to 5 where 0 = no significant change and 5 = whole organ or tissue affected for each observation. Tumor tissue, and other organs, for flow cytometry analyses were enzyme-digested to release single cells as previously described^41^. In brief, tissues were minced using scalpels, and then single cells were liberated by incubation for 60 mins at 37°C with 1 mg/ml Collagenase I from *Clostridium Histolyticum* (Sigma-Aldrich) and 0.1 mg/ml Deoxyribonuclease I (AppliChem) in RPMI (Gibco). Released cells were then passed through a 70 μm cell strainer prior to staining for flow cytometry analyses. Viable cells were numerated using a hemocytometer with trypan blue (Sigma-Aldrich) exclusion.

### Bioluminescence Imaging

For assessing Luc bio-distribution *in vivo* mice were injected intraperitoneally (i.p.) with 200μl (15mg/ml) XenoLight D-luciferin (PerkinElmer) in sterile PBS 10 mins prior to imaging to detect Click Beetle Luciferase. *Renilla reniformis* Luciferase was detected using 100μl RediJect Coelenterazine Bioluminescent Substrate (PrekinElmer) immediately prior to imaging. Animals were anesthetized for imaging and emitted light was detected using the *In vivo* Imaging System (IVIS^®^) Lumina Series III (PerkinElmer) and data analyzed using the Living Image software (PerkinElmer). Light was quantified in photons/second/unit area.

### Immunohistochemistry

To measure hypoxia within tissues, mice were injected i.p. with 60mg/kg pimonidazole HCl (Hypoxyprobe, HPI, Inc) dissolved in PBS, 2 h before sacrifice. Tissues were transferred into formalin solution (10%) neutral buffered (Sigma-Aldrich) for at least 24 h prior to paraffin embedding. Paraffin-embedded tissues were cut into 5 μm sections and mounted onto glass microscope slides (VWR). Sections were dewaxed in Histo-Clear™ (National Diagnostics) for 6 mins, prior to rehydration through 100%, 95%, 70% ethanol and tap water for 3 min each at room temp (RT). Antigens were retrieved using Access Revelation (Biocore LLC) at 95°C for 20 mins. Sections were washed three times in 100 mM Tris, 140 mM NaCl, 0.1% Tween 20 pH7.4 (TBST) wash buffer for 10 mins prior to applying a wax circle. Tissue peroxidases were quenched using 0.3% H_2_O_2_ (Sigma-Aldrich) for 10 mins at RT. Sections were washed again in TBST prior to blocking with 10% goat serum (Sigma-Aldrich) 0.1% Triton X-100 in TBST for 1 h at RT. The stable protein adducts formed with the reductively activated pimonidazole in hypoxic tissue were detected using rabbit anti-pimonidazole antisera (1:100 Pab2627, Hypoxyprobe, HPI, Inc) O.N. at 4°C. Sections were washed with TBST as above and bound rabbit IgG was detected using Dako EnVision™+ System-HRP (DAB) (K4010, Agilent Technologies) according to the manufacturer’s instructions. Tissue sections were counterstained with hematoxylin and washed clear with tap water prior to dehydration through 70%, 95%, 100% ethanol, and Histo-Clear™ (National Diagnostics) for 3 mins each prior to mounting with a coverslip using DePex (SERVA).

FFPE human HNSCC tumor tissues sections were deparaffinized and dual-antibody stained using a BenchMark ULTRA IHC/ISH system (Roche). Deparaffinized sections were pretreated with Cell Conditioning 1 (CC1) buffer (Roche) for 36 minutes and then incubated with mouse anti-human CD3ε (1:50 F7.2.38 Dako) for 32 mins at 37°C followed by amplification and detection using *ultraView* Universal DAB Detection Kit (Roche). Subsequently, tissues were incubated with Rabbit anti-human HIF1α (1:1000 EP1215Y, Abcam) for 32 mins at 37°C followed by amplification and detection using *ultra*View Universal Alkaline Phosphatase Red Detection Kit (Roche). Tissues were counterstained with hematoxylin and bluing reagent (Roche), dehydrated and mounted with a cover slip.

Images were acquired using a NanoZoomer Digital Slide Scanner (Hamamatsu) and IHC staining for stable protein adducts of hypoxyprobe and HIF1α were quantified using an H-score which represented the sum of the respective stain intensities from 0-3 (3 being highest), multiplied by the percent area that each intensity occupied across the tumor. Intraepithelial T-cells (IETs) were scored as low/absent if CD3^+^ cells were sparse or absent and ‘high’ if prevalent in the stromal regions surrounding the tumor. Tumor infiltrating lymphocytes (TILs) were scored as ‘present’ if there was >3 areas across the section where CD3^+^ cells could be found within the tumor tissue.

### Immunofluorescence

Sections from formalin fixed paraffin embedded (FFPE) human head and neck cancer, principally oral cavity (tongue) and tonsil, were de-paraffinized and antigen retrieved using a Ventana^®^ BenchMark ULTRA (Roche Tissue Diagnostics). Immunofluorescence was performed as previously described^38^. The following antibodies were used at 1:100 dilutions, mouse anti-CD3ε (F7.2.38, Dako) and Rabbit anti-HIF1α (EP12151, Abcam). Primary antibodies were detected using donkey IgG antibodies purchased from Thermo Fisher Scientific at 1:100: AlexaFluor^®^ 488 anti-mouse IgG and AlexaFluor^®^ 568 anti-rabbit IgG. Nuclei were stained using 1.25 μg/ml 4’,6-diamidino-2-phenylindole dihydrochloride (DAPI) (Thermo Fisher Scientific). Images were acquired using a Nikon Eclipse Ti-E Inverted Spinning Disk confocal microscope system and associated NIS Elements software. Colocalization of staining was evaluated using thresholding on the NIS Elements Software.

### Flow cytometry

Flow cytometry was performed as previously described^45^. The following antibodies were purchased from eBioscience and were used at 1 μg/ml unless stated otherwise: anti-human CD3ε Brilliant Violet 421™ (SK7; Biolegend^®^), anti-human CD8α Alexa Fluor 488 (RPA-T8), anti-human CD4 PE (RPA-T4), anti-human CD45 Brilliant Violet 510™ (HI30 Biolegend^®^), anti-mouse CD4 FITC (Clone: RM4-5), anti-mouse CD8α eFluor^^®^^450 (Clone: 53-6.7), antimouse CD3ε PE (Clone: 145-2C11), neutralizing anti-mouse CD16/CD32 (Clone: 2.4G2). Background fluorescence was established using fluorescence minus one staining. The T1E28z CAR was stained with a biotinylated anti-human EGF antibody (Bio-Techne: BAF236) and detected using Streptavidin APC. Human ErbB family members were detected using anti-ErbB1 (ICR62), anti-ErbB2 (ICR12; both ICR antibodies were gifts of Professor Suzanne Eccles, Institute of Cancer Research, Sutton), anti-ErbB3 PE (BioLegend^®^, 1B4C3), anti-ErbB4 (NOVUS Biologicals, Clone: H4.77.16). Antibodies that there not conjugated to a fluorochrome were detected using goat anti-Mouse IgG (H+L) highly cross-adsorbed secondary conjugated to Alexa Fluor Plus 488 (Thermo Fisher Scientific) or goat anti-rat IgG APC (BioLegend^®^) as appropriate. eGFP was detected by its native fluorescence. Dead cells and red blood cells were excluded using 1 μg/ml 7-amino actinomycin D (Cayman Chemical Company) alongside anti-Ter-119 PerCP-Cy5.5 (Ter-119; Thermo Fisher Scientific). Data were collected on a BD FACS Canto II (BD Biosciences). Data was analyzed using FlowJo software (Freestar Inc.).

### Computational analysis of cancer patient data

RSEM normalized expression datasets from the Cancer Genome Atlas (TCGA) were downloaded from the Broad Institute Firehose resource (https://gdac.broadinstitute.org/). The HRE-regulated gene expression signature was generated by taking the mean normalized log2-transformed expression value of the component signature genes. The HRE-regulated gene signature was comprised of HRE-regulated genes for which a positive correlation was observed between all genes. As *VEGFA* and *ENO1* did not correlate with each other, but were both correlated with the rest of the prospective signature candidates (Supplementary Fig 8a), we elected to include *VEGFA* over *ENO1* in the signature as, by comparison, *VEGFA*-inclusive signature provided a greater sensitivity for tumor T-stage and patient prognosis. This final signature included genes associated with glucose metabolism *(ALDOA, PGK1, SLC2A1),* pH regulation *(CA9)* and angiogenesis (*VEGFA*). All TCGA data was analyzed using R version 3.5.1 (https://www.r-project.org/). Overall survival analyses were generated by partitioning all HNSCC patients into quartiles based on hypoxia score and taking the top and bottom quartile expression ranked hypoxia score values. Kaplan-Meier survival curves were plotted using GraphPad Prism (GraphPad).

### Statistics

Normality and homogeneity of variance were determined using a Shapiro-Wilk normality test and an F-test respectively. Statistical significance was then determined using a two-sided unpaired Students *t* test for parametric or Mann-Whitney test for nonparametric data using GraphPad Prism 6 software. When comparing paired data, a paired ratio Students *t* test was performed. A Welch’s correction was applied when comparing groups with unequal variances. Statistical analysis of tumor growth curves was performed as described^46^. Correlation analyses were performed using Pearson correlation. The pairwise Wilcoxon Rank Sum Test with Benjamini-Hochberg correction was used to measure statistical differences between clinical groups and hypoxia score in cancer patient data from the TCGA. The log-rank (Mantel-Cox) test was used to determine statistical significance for overall survival in cancer patient data from TCGA. No outliers were excluded from any data presented.

### Study approval

The use of animals for this study was approved by the Ethical Review Committee at King’s College London and the Home Office, UK. Human HNSCC tumor tissue was obtained with informed consent under ethical approval from the King’s Health Partners Head and Neck Cancer Biobank (REC reference 12/EE/0493). Blood was obtained from healthy volunteers under approval of the Guy’s and St Thomas’ Research Ethics Committee (REC reference 09/H0804/92).

## Acknowledgements

Dr Yasmin Haque, King’s College London, for cell sorting and flow cytometry assistance. The authors would like to thank Dr Adam Ajina (KCL) for taking blood samples, Miss Dominika Sosnowska for help with tail vein injections and Dr Gilbert Fruhwirth for helpful discussion. This work was funded by the European Research Council (335326) and King’s Commercialisation Institute. P.K. and J.W.O. are supported by the UK Medical Research Council (MR/N013700/1) and are KCL members of the MRC Doctoral Training Partnership in Biomedical Sciences. The research was supported by the Cancer Research UK King’s Health Partners Centre and Experimental Cancer Medicine Centre at King’s College London, and the National Institute for Health Research (NIHR) Biomedical Research Centre based at Guy’s and St Thomas’ NHS Foundation Trust and King’s College London. The views expressed are those of the authors and not necessarily those of the NHS, the NIHR or the Department of Health.

## Authors contributions

P.K., J.M., J.N.A. conceived the project, designed the approach, interpreted the data and wrote the manuscript. P.K., J.W.O., K.I.L., R.H., C.L.S., M.O., M.Y.M.T, T.M., D.L.Y. performed experiments and interpreted the data. D.M.D., N.W., C.G., S.T. provided key expertise and interpretation.

## Data availability

Requests should be made to J.N.A.

## Extended data

**Extended Data Fig. 1.**
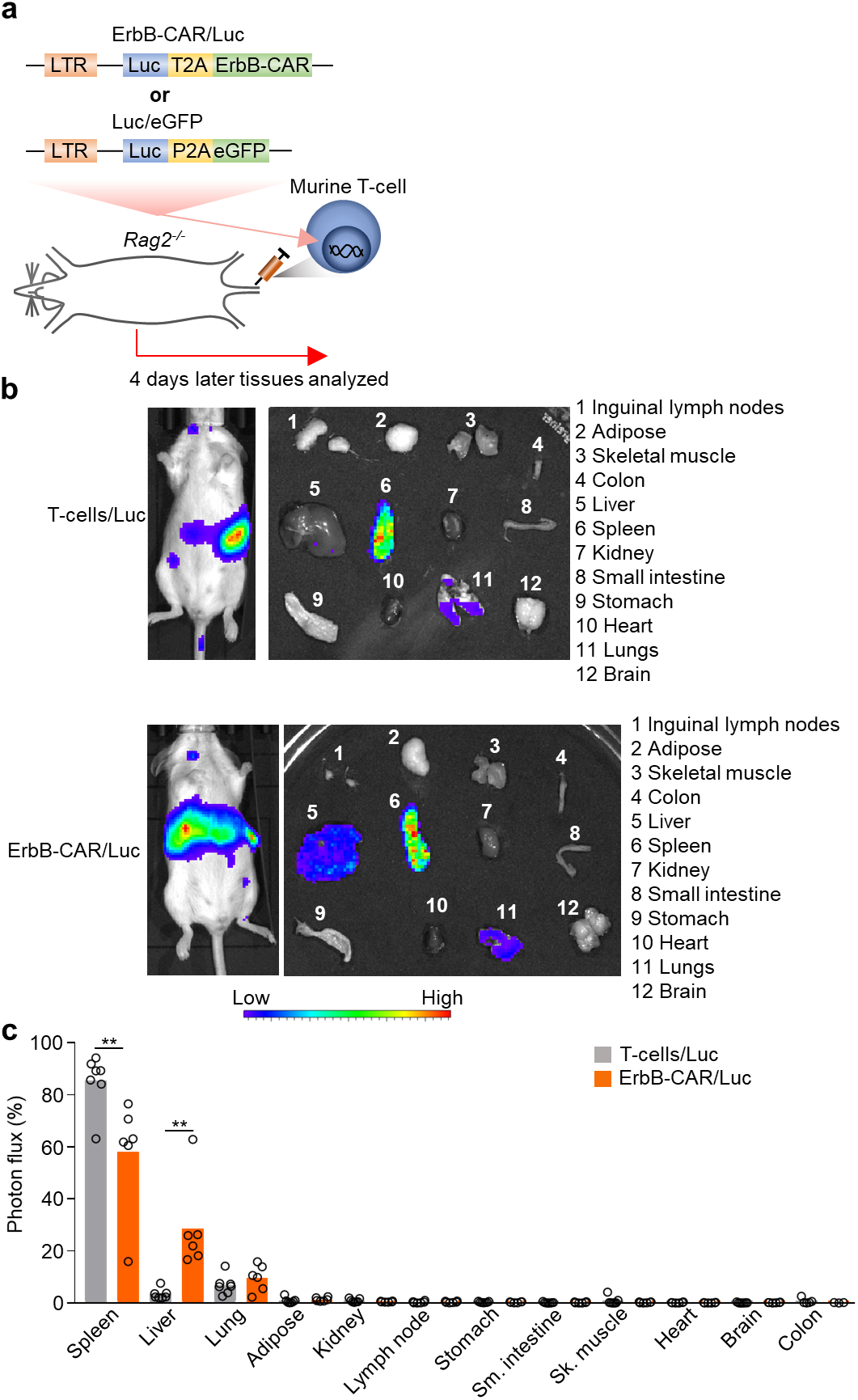
*In vivo* biodistribution of Luc-expressing murine T-cells and ErbB-CAR T-cells. (**a**) Schematic diagram depicting the Luc reporter constructs, and their modular arrangements, which were stably transduced into murine T-cells for the data in this Figure. (**b**) 5×10^6^ ErbB-CAR T-cells/Luc or control T-cells/Luc were infused into *Rag2^-/-^* mice and 4 days later imaged for bioluminescence. Shown are representative images of whole body (left) or post sacrifice dissected organs (right) emitted light. (**c**) Quantification of the photon flux from each organ/s/unit area (across n = 6 individual mice), displayed is the photons from each organ as a percent of all dissected organs. Bar charts show the group mean and each dot represents each individual mouse in the cohort. ** *P* < 0.01

**Extended Data Fig. 2.**
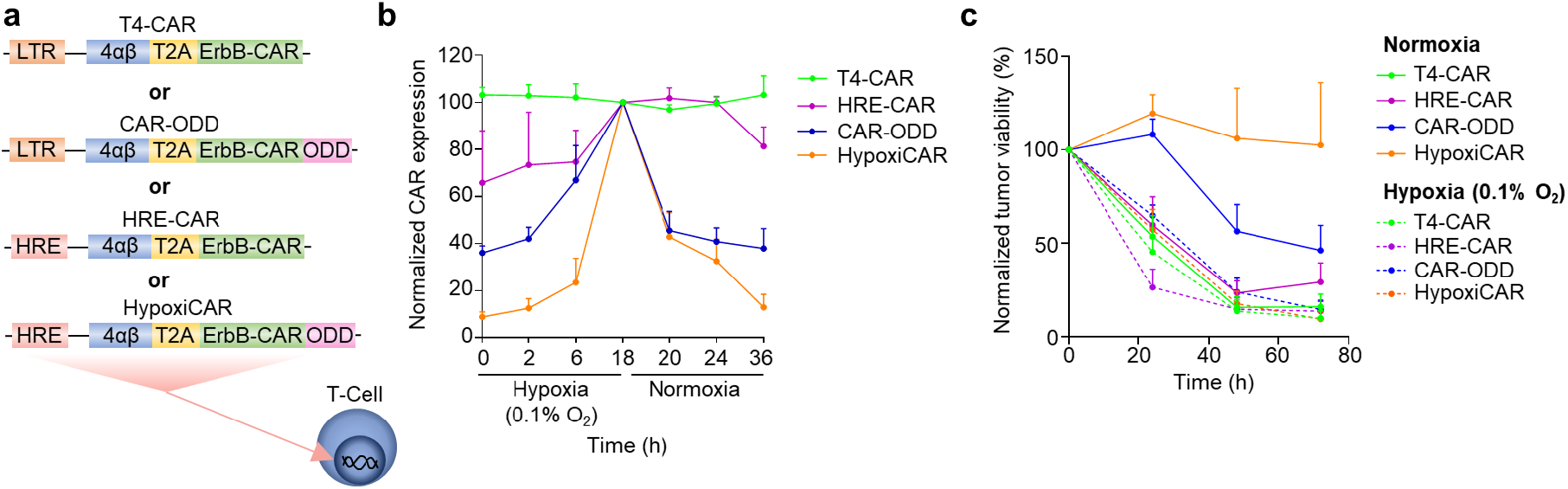
HypoxiCAR’s dual oxygen-sensing modules synergize to provide superior stringency in sensing hypoxia. (**a**) Schematic diagram depicting the constructs, and their modular arrangements, which were stably transduced into human T-cells for the data in this Figure. (**b**) Surface CAR expression on T4-CAR, HRE-CAR, CARODD and HypoxiCAR human T-cells at the indicated times under hypoxia (0.1% O_2_) or normoxia assessed using flow cytometry analyses. Values normalized to 18h hypoxia (n=4). (**c**) *In vitro* SKOV3 tumor cell killing by T4-CAR, HRE-CAR, CAR-ODD and HypoxiCAR human T-cells (CAR^+^ effector T-cell to target tumor cell ratio 4:1) in normoxic and hypoxic conditions (0.1% O_2_). Line charts, the dots mark mean and error bars s.e.m.

**Extended Data Fig. 3.**
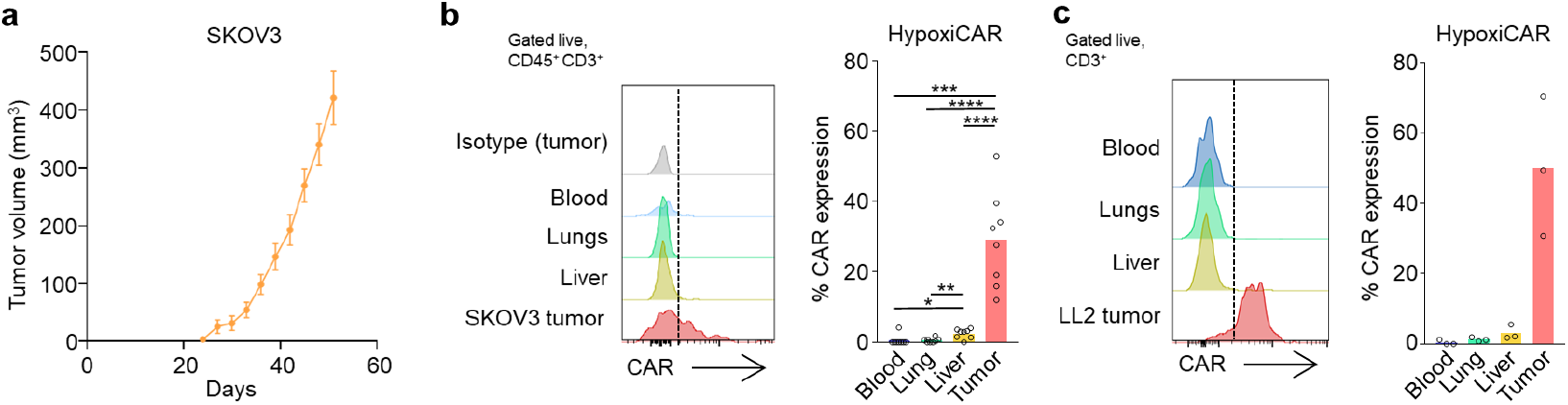
HypoxiCAR T-cells selectively express CAR in both SKOV3 and LL2 tumors. (**a**) Growth curve of SKOV3 tumors grown in NSG mice (n = 6 mice). Dots mark the mean and error bars the s.e.m. (**b**) Representative stacked histograms showing gated live CD45^+^ CD3^+^ events surface CAR expression on HypoxiCAR T-cells in the indicated enzyme-dispersed tissues and blood of a SKOV3 tumor-bearing mouse injected both i.v. and i.t. with 7.5×10^5^ and 2.5×10^5^ of human HypoxiCAR T-cells, respectively, 72 h prior to sacrifice. Histograms are from gated live nucleated cells (7AAD^-^, Ter119^-^), CD45^+^ CD3^+^ T-cells (as per supplementary Fig. 6b) alongside an isotype control stained tumor (grey histogram) (left) and the quantification of T-cells with detectable CAR in the respective tissues across n = 8 individual mice (right). (**c**) Equivalent experiment as described in (b), but where human HypoxiCAR T-cells were injected into LL2 tumor bearing *Rag2^-/-^* mice (across n = 3 individual mice). Bar charts show the group mean and each dot represents an individual healthy mouse. * *P* < 0.05, ** *P* < 0.01, *** *P* < 0.001, **** *P* < 0.0001.

**Extended Data Fig. 4.**
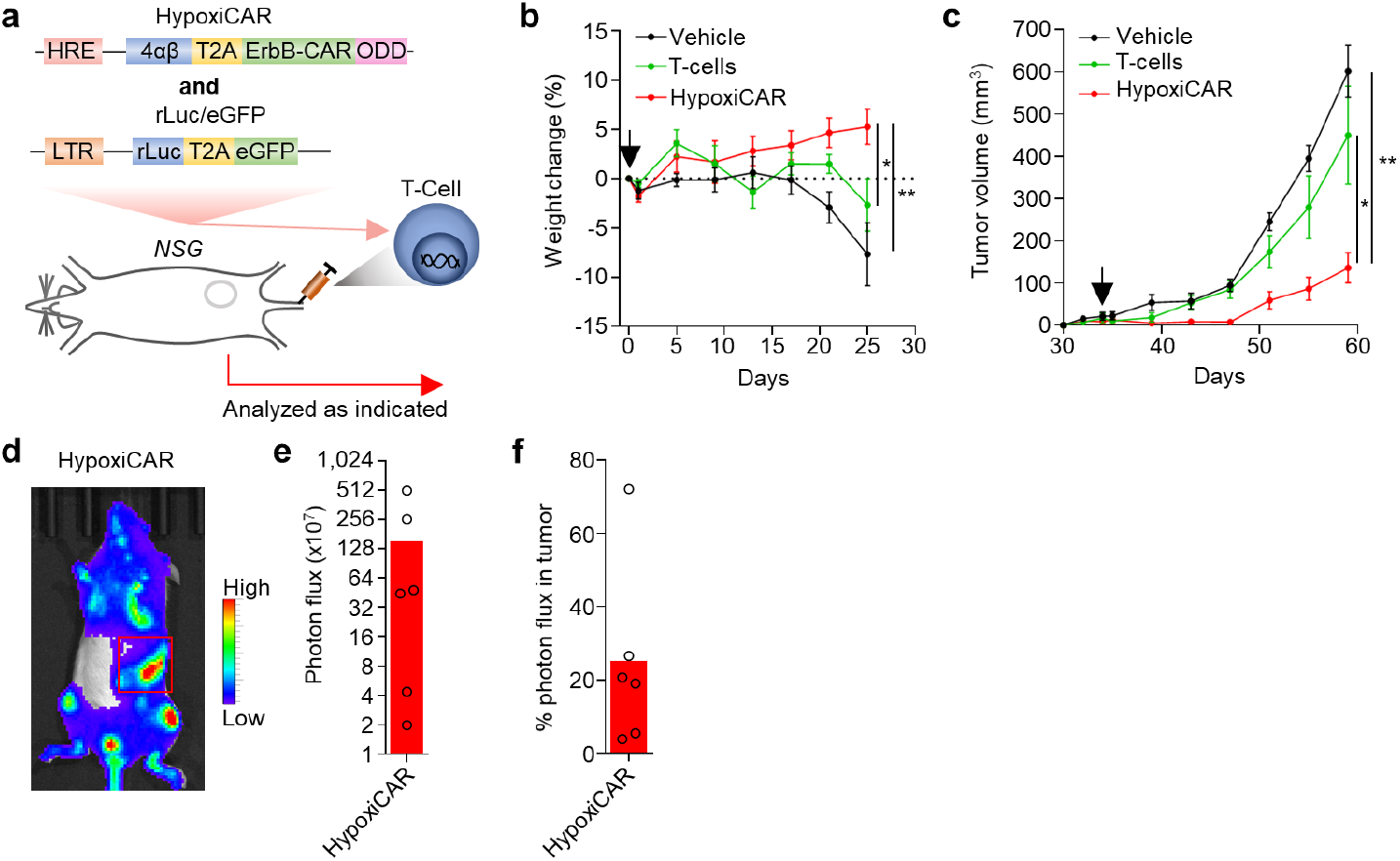
HypoxiCAR T-cells provide anti-tumor efficacy against established SKOV3 tumors. 32 days post subcutaneous SKOV3 tumor cell inoculation, when tumors were palpable, mice were infused i.v. with either vehicle (n=6) or 10×10^6^ reporter HypoxiCAR (n=7) or non-transduced T-cells (control, n=5). (**a**) Schematic diagram depicting the experiment in which HypoxiCAR T-cells were co-transduced to concurrently express constitutive rLuc/eGFP. (**b**) Weight change of the mice shown as days post T-cell infusion. (**c**) SKOV3 tumor growth curves shown as days post tumor cell inoculation. (**d**) Bioluminescence imaging was performed on the whole body of mice to track the biodistribution of the infused HypoxiCAR T-cells at day 26 post infusion (day 60 post tumor cell inoculation). Red box marks the tumor. (**e-f**) Quantification of the photons/s/unit area (photon flux) across the whole body (**e**) and the % photon flux signal detected specifically in the tumor (**f**). Arrow marks the point of CAR T-cell infusion. All experiments are representative of a biological repeat. Bar charts shows the mean and each dot an individual mouse. In line charts, the dots mark the mean and error bars s.e.m. * *P*<0.05, ** *P*<0.01.

