## Supplemental Material for "Hypoxia-Sensing CAR T-Cells Provide Safety and Efficacy in Treating Solid Tumors"

### Supplementary information

| Tissue | Pathology | Mouse | | | | | |
|---|---|---|---|---|---|---|---|
| | | 1 | 2 | 3 | 4 | 5 | 6 |
| | | Vehicle | | | | | |
| Lung | | 0 | 0 | 0 | 0 | 0 | 0 |
| Liver | Inflammatory cell infiltrate, parenchymal | 0 | 1 | 0 | 0 | 0 | 0 |
| | Inflammatory cell infiltrate, periportal | 0 | 1 | 0 | 0 | 0 | 0 |
| | Increased mitoses | 0 | 1 | 0 | 1 | 0 | 0 |
| | Pigmented macrophages | 0 | 1 | 0 | 0 | 0 | 0 |
| Kidney | | 0 | 0 | 0 | 0 | 0 | 0 |
| | | T-cells | | | | | |
| Lung | | 0 | 0 | 0 | 0 | 0 | 0 |
| Liver | | 0 | 0 | 0 | 0 | 0 | 0 |
| Kidney | | 0 | 0 | 0 | 0 | 0 | 0 |
| | | ErbB-CAR T-cells | | | | | |
| Lung | Inflammatory cell infiltrate, vascular | 3 | 3 | 3 | 2 | 3 | 0 |
| | Increased cellularity, alveolar septae | 1 | 2 | 1 | 2 | 1 | 0 |
| Liver | Single cell necrosis/apoptosis, scattered | 1 | 0 | 1 | 1 | 0 | 0 |
| | Inflammatory cell infiltrate, parenchymal | 1 | 1 | 1 | 1 | 0 | 1 |
| | Inflammatory cell infiltrate, perivascular | 2 | 2 | 1 | 2 | 2 | 0 |
| | Increased mitoses | 1 | 0 | 0 | 1 | 0 | 0 |
| Kidney | Cyst, cortex | 0 | P | 0 | 0 | 0 | 0 |
| | Tubular basophilia | 0 | 0 | 0 | 0 | 1 | 0 |
| | Inflammatory cell infiltrate, subcapsular | 0 | 1 | 0 | 0 | 0 | 0 |

**Supplementary Table 1│Histopathological evaluation of ErbB-targeted CAR T-cell mediated inflammation in healthy organs.** H&E stained tissue sections from each mouse and tissue were scored blinded. NSG mice were infused i.v. with $4.5x10^6$ T4-CAR T-cells or control (non-transduced) human T-cells, or vehicle and the indicated tissues were harvested at day 5 post-infusion for histological examination. The table presents all identified histological observations for each group and tissue. Observations in vehicle treated cohort and in the T4-CAR T-cell cohort were consistent with background spontaneous findings in most mice strains. Scores are a non-linear, semi-quantitative grading system from 0 to 5 where 0 = no significant change and 5 = whole organ or tissue affected for each observation[47]. For some observations grading was not appropriate and they are scored ‘P’ for present.

| Tissue | Pathology | Mouse | | | | | |
|---|---|---|---|---|---|---|---|
| | | 1 | 2 | 3 | 4 | 5 | 6 |
| | | | | Vehicle | | | |
| Lung | | 0 | 0 | 0 | 0 | 0 | 0 |
| Liver | Inflammatory cell infiltrate, parenchymal | 0 | 1 | 0 | 0 | 0 | 0 |
| | Inflammatory cell infiltrate, periportal | 0 | 2 | 0 | 0 | 0 | 0 |
| | Focal necrosis | 0 | 2 | 0 | 0 | 0 | 0 |
| | Increased mitoses | 0 | 0 | 0 | 1 | 0 | 1 |
| Kidney | Tubular basophilia | 0 | 0 | 1 | 0 | 0 | 0 |
| | | | | T-cells | | | |
| Lung | | 0 | 0 | 0 | 0 | 0 | - |
| Liver | Increased mitoses | 0 | 0 | 1 | 0 | 0 | - |
| Kidney | | 0 | 0 | 0 | 0 | 0 | - |
| | | | | T4-CAR T-cells | | | |
| Lung | Inflammatory cell infiltrate, vascular | 3 | 3 | 3 | 3 | 3 | - |
| | Increased cellularity, alveolar septae | 2 | 2 | 2 | 2 | 2 | - |
| Liver | Single cell necrosis/apoptosis, scattered | 0 | 1 | 1 | 1 | 1 | - |
| | Inflammatory cell infiltrate, parenchymal | 1 | 1 | 1 | 1 | 1 | - |
| | Inflammatory cell infiltrate, perivascular | 2 | 3 | 2 | 2 | 1 | - |
| | Increased mitoses | 0 | 0 | 0 | 1 | 1 | - |
| Kidney | Inflammatory cell infiltrate, subcapsular | 0 | 1 | 0 | 0 | 0 | - |
| | Nephropathy | 0 | 0 | 0 | 3 | 0 | - |
| | | | | HypoxiCAR T-cells | | | |
| Lung | | 0 | 0 | 0 | 0 | 0 | 0 |
| Liver | Increased mitoses | 1 | 0 | 1 | 1 | 0 | 0 |
| Kidney | | 0 | 0 | 0 | 0 | 0 | 0 |

**Supplementary Table 2 Pathohistological evaluation of organs from mice infused with either T4-CAR or HypoxiCAR T-cells.** H&E stained tissue sections from each mouse and tissue were scored blinded. NSG mice were infused i.v. with $4.5x10^6$ T4-CAR, HypoxiCAR, control (non-transduced) human T-cells or vehicle and the indicated tissues were harvested at day 5 post infusion for histological examination. The table presents all identified histological observations. Observations in vehicle treated cohort were consistent with background spontaneous findings in most mice strains. Scores are a non-linear, semi-quantitative grading system from 0 to 5, where 0 = no significant change and 5 = whole organ or tissue affected for each observation[47].

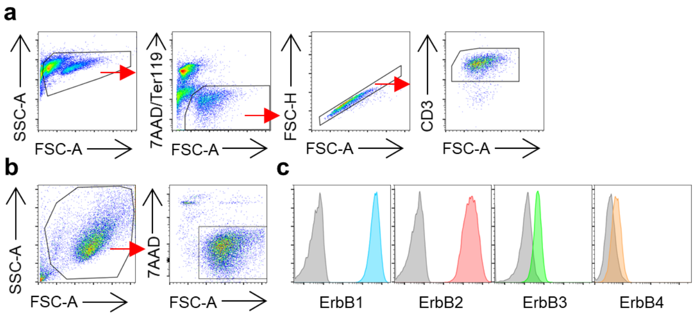


**Supplementary Fig. 1 | Flow cytometry gating strategies and tumor cell ErbB receptor expression**. (**a**) Example of the flow cytometry gating strategy for live ($7AAD^-$), singlet, $CD3^+$ T-cells. (**b**) Flow cytometry dot plots showing the gating strategy for live ($7AAD^-$) HN3 cells. (**c**) Live gated HN3 cells surface expression of ErbB1-4 (colored histograms) against their respective isotype control staining (grey histograms).

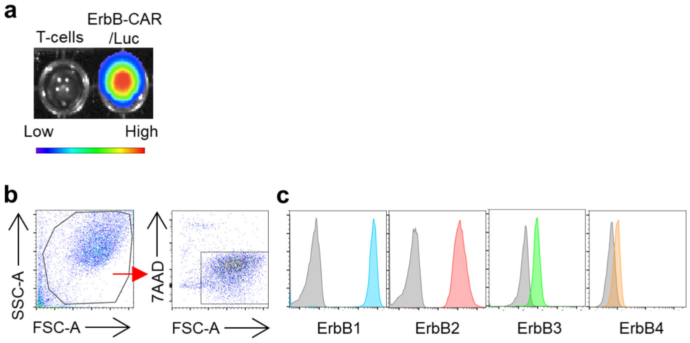


**Supplementary Fig. 2│Validation of the ErbB-CAR reporter T-cells and ErbB receptor expression on SKOV3 cells**. (**a**) *In vitro* validation of functional Luc expression by the human ErbB-CAR reporter T-cells versus non-transduced T-cells prior to infusion into mice, through detection of emitted light upon exposure to luciferin substrate in wells of a 96-well plate. (**b**) Flow cytometry dot plots showing the gating strategy for live (7AAD$^{-}$) SKOV3 cells. (**c**) Live gated SKOV3 cells surface expression of ErbB1-4 (colored histograms) against their respective isotype control staining (grey histograms).

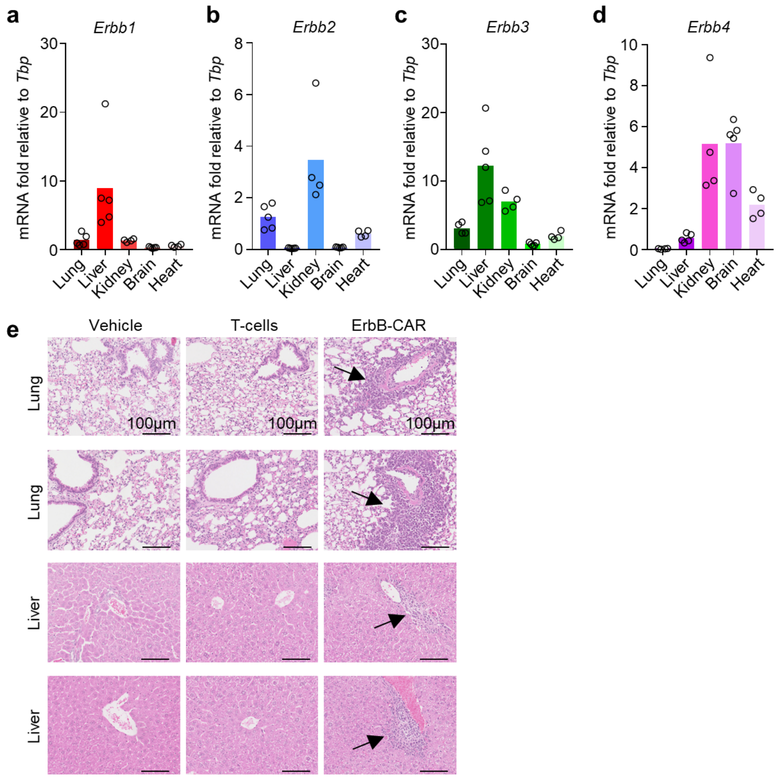


**Supplementary Fig. 3│ErbB1-4 expression in vital organs and ErbB-CAR T-cell mediated inflammation in the lungs and liver post i.v. infusion.** (**a-d**) mRNA expression of *Erbb1* (**a**), *Erbb2* (**b**), *Erbb3* (**c**) *Erbb4* (**d**) genes relative to the housekeeping gene *Tbp* in the indicated tissues (n = 6). (**e**) Additional H&E stained sections from the indicated tissue 5 days post infusion with 4.5x10$^{6}$ T4-CAR, non-transduced human T-cells or vehicle control. Arrows indicated myeloid infiltration zones as a surrogate marker of inflammation. Scale bar denotes 100µm. Bar charts show the group mean and each dot represents each individual mouse.

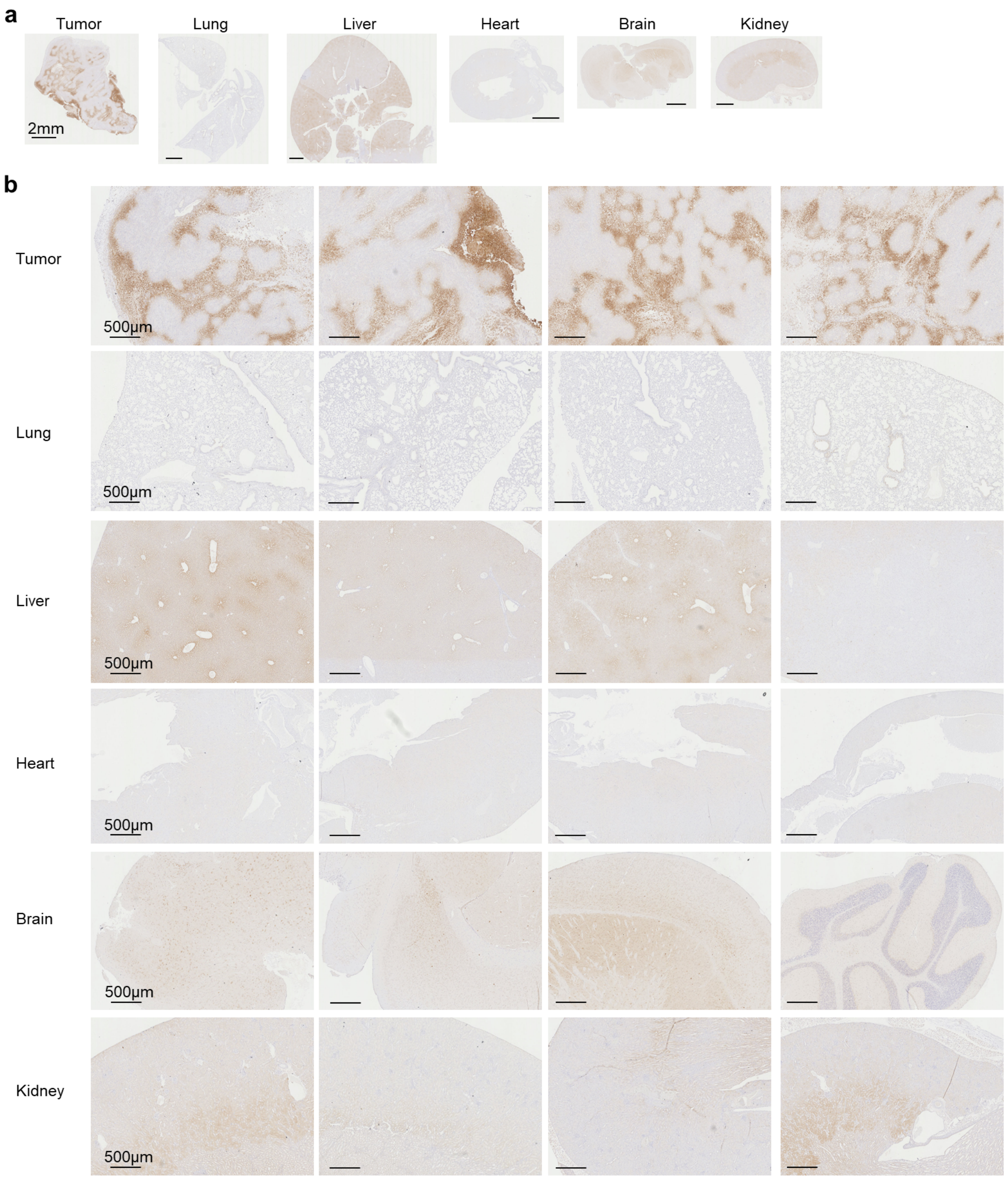


**Supplementary Fig. 4│Hypoxia differentiates the tumor microenvironment**. Additional images assessing hypoxia, detected in tissue sections by immunoperoxidase staining of stable protein adducts formed with reductively-activated pimonidazole in HN3 tumors and healthy organs from tumor bearing NSG mice. (**a**) Representative whole tissue images (**b**) Representative regions from the individual tissues (rows). Images randomly selected across a cohort of n = 5 individual mice.

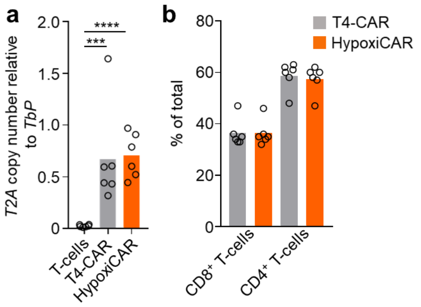


**Supplementary Fig. 5│Characterization of CAR T-cell** (**a**) Genomic DNA from T4-CAR, HypoxiCAR and non-transduced human T-cell preparations subjected to qPCR for *T2A* copy number (retroviral integration) relative to that of *Tbp* (endogenous gene) in the genomic DNA. (**b**) The relative prevalence of CD4$^+$ and CD8$^+$ T-cells assessed using flow cytometry in the T-cells, T4-CAR and HypoxiCAR preparations**.** (n = 6 individual healthy donors). Bar charts show the group mean and each dot represents an individual healthy donor in the group. *** $P < 0.001$, **** $P < 0.0001$.

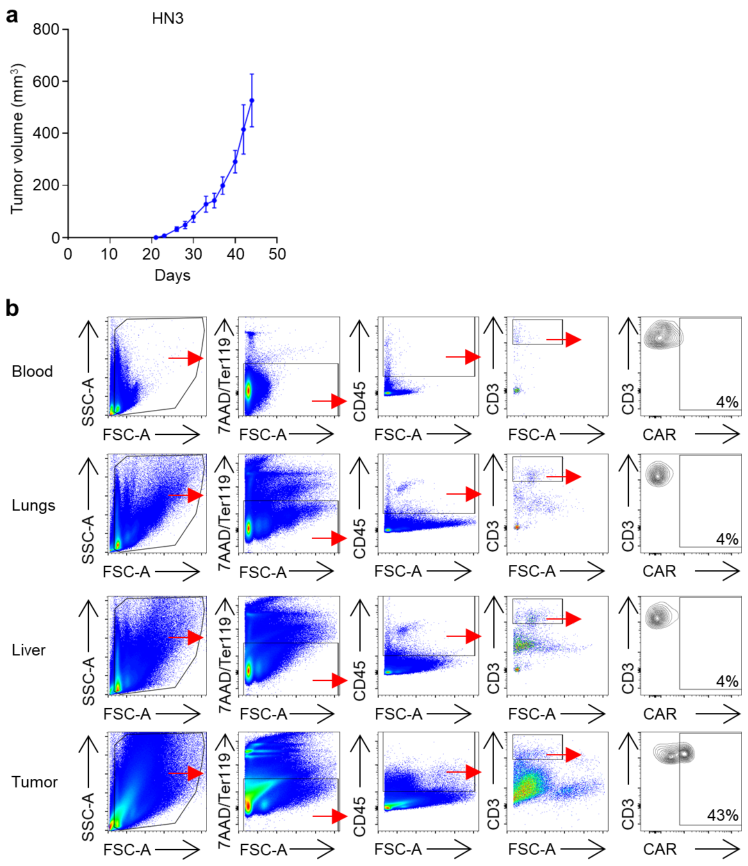


**Supplementary Fig. 6│HypoxiCAR T-cells selectively express CAR in the hypoxic tumor microenvironment**. (**a**) Growth curve of HN3 tumors grown in NSG mice (n = 6 mice). Dots mark the mean and error bars the s.e.m. (**b**) Representative flow cytometry dot/contour plots for the gating strategy from enzyme-dispersed tumors and healthy tissues alongside the blood from mice which had been injected both i.v. and i.t. with $7.5x10^5$ and $2.5x10^5$ of human HypoxiCAR T-cells, respectively, 72 h prior to sacrifice. Stained for live cells (7AAD$^-$, Ter119$^-$), CD45$^+$ CD3$^+$ T-cells and their surface CAR expression. Positive gates were applied based on isotype staining.

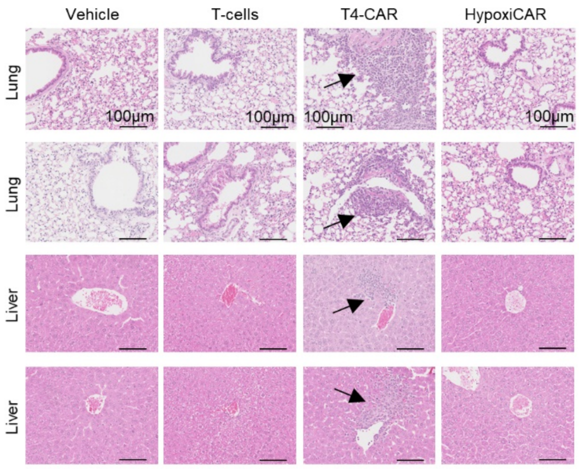


**Supplementary Fig. 7│HypoxiCAR T-cells do not evoke inflammation in the lungs and liver.** Additional H&E stained sections from each indicated tissue from mice 5 days post i.v. infusion with 4.5x$10^6$ T4-CAR, HypoxiCAR, non-transduced human T-cells or vehicle treated animals. Arrows indicated myeloid infiltration zones as a surrogate marker of inflammation. Scale bar denotes 100µm.

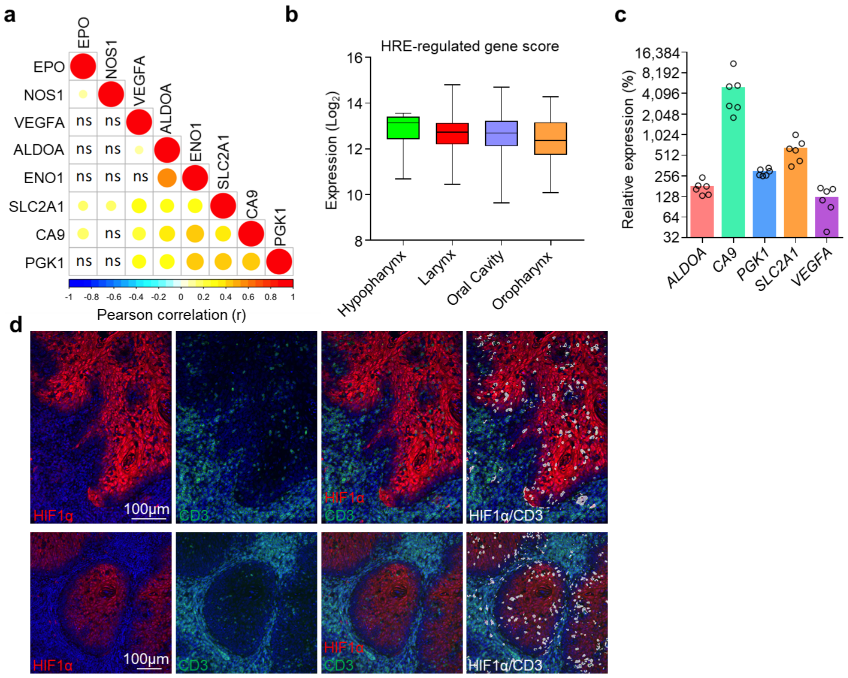


**Supplementary Fig. 8│HRE-regulated gene signature and HIF1α in HNSCCs.** (**a**) Correlation plot showing pairwise correlation of HRE-regulated genes in the HNSCC. The size of the dot represents the *P* value of the correlation where *P* > 0.05 and the color of the dot represents the Pearson correlation coefficient (r). (**b**) Expression of the HRE-regulated gene score in HNSCC based on subtype (hypopharynx n = 10, larynx n = 116, oral cavity n = 316, oropharynx n = 79). (**c**) mRNA expression of the indicated HRE-regulated genes relative to the housekeeping gene *TBP* in 500mm$^3$ SKOV3 tumors grown in NSG mice normalized to the respective gene expression of the *in vitro* cultured cells (n = 6). (**d**) Additional representative confocal images from sections from two further HNSCCs stained with DAPI (nuclei; blue) and antibodies against CD3 (green) and HIF1α (red), white events denote CD3 and HIF1α co-localizing pixels. Box plots show median and upper/lower quartiles, whiskers show highest and lowest value. Bar chart shows the group mean and each dot represents an individual mouse and tumor.
